# eEF2 kinase enhances the expression of PD-L1 by promoting the translation of its mRNA

**DOI:** 10.1101/2020.09.01.278655

**Authors:** Yu Wu, Jianling Xie, Xin Jin, Roman V. Lenchine, Xuemin Wang, Danielle M. Fang, Zeyad D. Nassar, Lisa M. Butler, Jing Li, Christopher G. Proud

**Author notes:** these authors contributed equally to this study (Yu Wu, Jianling Xie). Corresponding authors: Jing Li: Key Laboratory of Marine Drugs, Chinese Ministry of Education, School of Medicine and Pharmacy, Ocean University of China, Yushan Road, Qingdao, China., Christopher G. Proud: Lifelong Health, South Australian Health and Medical Research Institute, North Terrace, Adelaide, SA5000, Australia.

## Abstract

Emerging advances in cancer therapy have transformed the landscape from conventional therapies towards cancer immunotherapy regimens. Recent discoveries have resulted in the development of clinical immune checkpoint inhibitors that are ‘game-changers’ for cancer immunotherapy. Here we show that eEF2K, an atypical protein kinase that inhibits the elongation stage of protein synthesis, actually promotes the synthesis of PD-L1, an immune checkpoint protein which helps cancer cells to escape from immunosurveillance. Ablation of eEF2K in prostate and lung cancer cells markedly reduced the expression levels of the PD-L1 protein. We show that eEF2K promotes the association of *PD-L1* mRNAs with translationally active polyribosomes and that translation of the *PD-L1* mRNA is regulated by a uORF (upstream open reading-frame) within its 5’-UTR (5’-untranslated region) which starts with a non-canonical CUG codon. This inhibitory effect is attenuated by eEF2K thereby allowing higher levels of translation of the PD-L1 coding region and enhanced expression of the PD-L1 protein. Moreover, eEF2K-depleted cancer cells are more vulnerable to immune attack by natural killer cells. Therefore, control of translation elongation can modulate the translation of this specific mRNA, one which contains an uORF that starts with CUG, and perhaps others that contain a similar feature. Taken together, our data reveal that eEF2K regulates PD-L1 expression at the level of the translation of its mRNA by virtue of a uORF in its 5’-region. This, and other roles of eEF2K in cancer cell biology (e.g., in cell survival and migration), may be exploited for the design of future therapeutic strategies.

## Introduction

It is now clear that immunotherapy is a promising and effective approach to tackling a range of cancers (reviewed [1]). However, cancer cells can resist immune attack through so-called checkpoints, the two most-widely studied of which are programmed cell death protein 1 (PD-1) and cytotoxic T-lymphocyte protein 4) [1]. PD-1, which is expressed on the surface of T lymphocytes, binds ligands expressed on other cells which, through PD-1, repress the activity of T cells. One such ligand is the protein PD-ligand 1 (PD-L1; [1], also termed CD274) which can be induced by signals such as interferon-γ (IFNγ), a cytokine. It is increasingly clear that countering these so-called immune check-points promotes immune responses, including against cancer cells. Indeed, antibodies that block either PD-1 or PD-L1 have shown striking efficacy in tackling a number of cancers (see, e.g., [1]).

As such, it is important to understand the mechanisms that control the expression of PD-L1 on cancer cells and thus enable them to ‘hide’ from immune surveillance. The expression of PD-L1 is known to be controlled at multiple levels including transcription of its gene, the stability of its mRNA and by microRNAs [1, 2].

mRNA translation is subject to control by a range of sophisticated mechanisms downstream of multiple signalling pathways [3]. One such mechanism involves the modulation of the rate of the elongation phase of protein synthesis through the phosphorylation of eukaryotic elongation factor 2 (eEF2) by its cognate kinase, eEF2K. eEF2K is an atypical, Ca^2+^-dependent protein kinase which is activated under diverse stress conditions and is turned off by signalling through the anabolic mechanistic target of rapamycin complex 1 (mTORC1) signaling pathway [4]. Importantly, eEF2K is not required for mammalian viability, fertility or health under normal vivarium conditions [5], meaning that agents that block it activity are not expected to exert adverse on-target effects.

Interestingly, eEF2K plays role in tumour cell survival during nutrient depletion, probably by slowing down protein synthesis and thereby reducing the consumption of energy and amino acids [6-8] and in cancer cell migration and angiogenesis, by regulating the synthesis of proteins involved this process including some integrins [9, 10]. Indeed, a number of studies have shown that eEF2K and/or alteration of translation elongation rates affects the translation of specific mRNAs and thus the synthesis of individual types of proteins (e.g., [9, 11, 12]), although the mechanisms underlying this have remained unclear.

Here we show, for the first time, that the expression of the PD-L1 protein is promoted by eEF2K, and thus the control of the elongation phase of protein synthesis. We show that this involves a novel mechanism whereby eEF2K, suppresses the effect of an inhibitory upstream open reading-frame (uORF). This raises the possibility that other members of the set of mRNAs that contain such features [13] may also be controlled by eEF2K. Other, previously identified uORF-regulated mRNAs are controlled in a quite different way, via the translation initiation factors eIF2 and eIF2B [14].

These findings further strengthen the rationale that eEF2K is a potentially valuable target for cancer therapy, reflecting its role in processes including immune surveillance, as well as cell survival and cell migration.

## Materials and Methods

### Chemicals and reagents

All chemicals were from Merck (Frenchs Forest, NSW, Australia) unless otherwise specified. Human interferon γ (IFNγ), rapamycin and eFT508 were purchased from Jomar Life Research (Scoresby, Australia).

### Cell culture, treatment and lysis

Cell line authentications were performed by Garvan Molecular Genetics (Darlinghurst, NSW, Australia) and CellBank Australia (Westmead, NSW, Australia) for A549 and PC3 cells, respectively. Cells were routinely tested against mycoplasma contamination. PC3 (human prostate cancer) cells were maintained in Roswell Park Memorial Institute-1640 (RPMI-1640) media (2 g/L glucose) containing 10% (v/v) foetal bovine serum (FBS) and 1% penicillin/streptomycin (growth medium). Human lung carcinoma A549 cells expressing an inducible short hairpin RNA (shRNA) against eEF2K were generously provided by Janssen Pharmaceutica (Beerse, Belgium) [15]. To induce the knockdown of eEF2K, A549 cells were cultured with 1 µM isopropyl-β-D-1thiogalactopyranoside (IPTG) for 5 days prior to use. A549 cells were maintained in Dulbecco’s modified Eagle medium media (4.5 g/L glucose) containing 10% (v/v) FBS and 1% penicillin/streptomycin. NK-92 cells were cultured in α-MEM media containing 12.5% (v/v) FBS, 12.5% horse serum and 100 IU/ml IL-2. All cells were cultured at 37°C in 5% CO_2_ and 95% air. For pH-buffered media, pH was adjusted by adding NaHCO_3_.

After treatment, cells were lysed by scraping into ice-cold lysis buffer containing 1% (v/v) Triton X-100, 20 mM Tris-HCl pH 7.5, 150 mM NaCl, 1 mM EDTA, 1 mM EGTA, 2.5 mM Na_2_H_2_P_2_O_7_, 1 mM β-glycerophosphate, 1 mM Na_3_VO_4_, 1 mM dithiothreitol and protease inhibitor cocktail, unless otherwise stated. Lysates were spun at 4 °C, 16,000 x *g* for 10 min, the supernatants were kept and total protein concentrations were quantified by Bradford assay (Bio-Rad, Gladesville, NSW, Australia) following the manufacturer’s instructions. Normalized lysates were either kept at -20°C or subjected to further analysis.

### Generation of CRISPR-directed eEF2K^-/-^ cells

The eEF2K-KO CRISPR targeting vector was the GeneArt CD4 CRISPR Nuclease Vector (ThermoFisher Scientific, Scoresby, VIC, Australia). The guide ssDNA sequence was 50-AGTGAGCGGTATAGCTCCAG-30. PC3 cells were transfected with the CRISPR vector by nucleofection (Lonza, Mt Waverley, Australia). After 72 h, CD4-positive cells were enriched using magnetic CD4 Dynabeads (ThermoFisher Scientific) and then sorted into individual wells of 96-well plates using a BD FACSFusion flow cytometer (Becton, Dickinson & Company, Wayville, SA, Australia). Positive clones were selected by immunoblotting analysis for the absence of eEF2K and P-eEF2, and were further confirmed by Sanger sequencing analysis (Supplementary Figure S1; performed by the Australian Genome Research Facility, Adelaide, Australia).

### SDS-PAGE/WB analysis

Sodium dodecyl sulphate-polyacrylamide gel electrophoresis (SDS-PAGE) and Western blot (WB) analysis were performed as previously described [16]. Primary antibodies used were: New England Biolabs (NEB), Hitchin, Herts, UK: PD-L1 (catalog number 13684), eEF2 (2332), eIF4E (2067), ThermoFisher Scientific: P-eIF4E Ser209 (44-528-G); Sigma-Aldrich: β-actin (A5316); Eurogentec, Seraing, Belgium: P-eEF2 Thr56 and eEF2K (custom-made); Servicebio, Wuhan, Hubei, China: GAPDH (GB12002). Fluorescently tagged secondary antibodies were from ThermoFisher Scientific. Blots were scanned using a LI-COR Odyssey imaging system (Millenium Science, Mulgrave, Victoria, Australia). Blots were quantified using the LI-COR Image Studio Lite 4.0 software (LI-COR, Lincoln, NE, USA).

### Plasmid and transfection

pICtest2 vector encoding StaCFluc with different start codons (AUG or CUG) were described previously [17]. PD-L1 5’-UTR was introduced to the pICtest2 StaCFluc vector using the HindIII restriction site. Point mutations were introduced by PCR mutagenesis using the Pfu DNA polymerase (Promega, Alexandria, VIC, Australia). Cells were transfected using Lipofectamine® 3000 (Thermofisher Scientific, Adelaide, SA, Australia) according to manufacturer’s instructions.

### Polysome analysis

Polysome analysis was performed as described previously [18]. Briefly, 80% confluent cells grown on 10 cm diameter culture dishes were lysed with 300 μl of lysis buffer [10 mM NaCl, 10 mM MgCl_2_, 10 mM Tris–HCl, pH 7.5, 1% (v/v) Triton X-100, 1% sodium deoxycholate, 36 U/ml RNase inhibitor, 1 mM dithiothreitol] and layered onto a 20–50% (w/v) sucrose gradient containing 30 mM Tris–HCl, pH 7.5, 100 mM NaCl and 10 mM MgCl_2_, and centrifuged in a Beckman SW41 rotor for 150 min at 234,000 x *g*. Fractions were collected while monitoring absorbance at 254 nm. For RNA extraction, 1% (w/v) SDS and 0.15 mg/ml proteinase K were added to each fractions, 1:3 (v/v) phenol:chloroform, pH 4.5, was then added to the samples to extract RNA, and RNA was precipitated from the aqueous phase by the addition of 70% (v/v) isopropanol. RNA pellets were washed once with 80% (v/v) ethanol before dissolving in RNase/DNase-free water for further analysis. For qPCR analysis of sucrose gradient fractions, 1.2-kb kanamycin RNA provided by the reverse transcription kit, used as an internal control for the reverse transcription (RT) reaction, was added to the RNA samples isolated from each of the polysome fractions just before the RT.

### Real-time quantitative RT-PCR (qPCR) amplification analysis

Total RNA was extracted using TRIzol (ThermoFisher Scientific). cDNA was produced using the QuantiNova reverse transcription kit (Qiagen, Chadstone, VIC, Australia). For qPCR analysis of sucrose gradient fractions, 1.2-kb kanamycin RNA (Promega) was used as an internal control. qPCR was performed using the following human primers (5’–3’): *PD-L1*: forward: TGGCATTTGCTGAACGCATTT; reverse: TGCAGCCAGGTCTAATTGTTTT; *B2M*: forward: TGGGTTTCATCCATCCGACA, reverse: ACGGCAGGCATACTCATCTT. Samples were analyzed with PowerUp SYBR Green master mix (ThermoFisher Scientific) on an ABI Step One Plus qPCR instrument (Applied Biosystems, Cheshire, UK). For total RNA analysis, *ACTB* (Forward: CATGTACGTTGCTATCCAGGC; Reverse: CTCCTTAATGTCACGCACGAT) was used as the normalization control. The comparative threshold cycle (C_T_) method was applied to quantify mRNA levels.

### Luciferase assays

*Firefly* luciferase (Fluc) activity was measured with luciferase reporter assay systems (Promega) on a GloMax-discover multimode detection system (Promega) following the manufacturer’s instructions.

### Flow cytometry analysis

Following treatment, cells were trypsinized and incubated with PD-L1 antibody [NEB, catalog number 86744; 1:100 diluted in 0.1% BSA (w/v) in phosphate buffered saline (PBS)] for 30 min at 4 °C, cells were washed thrice with PBS and then incubated with Alexa Fluor-conjugated rabbit secondary antibody (Abcam, Melbourne, VIC, Australia) for 30 min. They were washed again thrice with PBS, the intensity of fluorescence signals was recorded using FACS CantoTM II flow cytometry [Becton Dickinson (BD) Biosciences, North Ryde, NSW, Australia], data were analyzed using FlowJo software version 10.2 (BD Biosciences).

### Lentiviral shRNA infection and NK-92 killing assays

Lentiviral non-targeting control (sh-NC) shRNA or shRNA targeting eEF2K (sh-eEF2K; sc-39011-V) were purchased from Santa Cruz Biotechnology. Cells were infected with sh-NC, sh-eEF2K, positive clones were screened by puromycin (8 μg/ml) selection. Cells were transfected with an empty vector (E.V.) or FLAG-eEF2K [19] where indicated. 48 h later, 3,500 cells per sample were seeded on 48-well plates overnight, NK-92 cells were then added to cancer cells at a 5:1 ratio for 3 (for PC3) or 8 (for A549) h. NK-92 cells and cell debris were removed by washing with PBS, and living cancer cells were then quantified by spectrometric analysis at OD (570 nm), followed by crystal violet staining [20]. Before staining, we also took microscopic images of the cells with a Cytation 5 multimode Reader (BioTek, Beijing, China). PD-1/PD-L1 Inhibitor 3 (Selleck 3S8158) was used as a positive control.

### Cytokine assays

To assess levels of granzyme B, IFN-γ and perforin in culture, 1 x 10^4^ PC3 cells per sample were seeded on 96-well plates. After 16 h, NK-92 cells were added to PC3 cells at a 5:1 ratio and cells were further for a further 12 h. Cytokines from culture media were measure using the following ELISA kits (Dakewe Biotech, Shenzhen, China) according to the manufacturer’s instructions: Granzyme B (1118502), Perforin (1118302), and IFN-γ (1110002).

### Gene expression and survival analysis from publicly available data

*EEF2K* and *CD274* (PD-L1) gene expression and patient survival data were retrieved from Cancer Genome Atlas (TCGA) and GEO (database: GSE56288) [21, 22]. Kaplan–Meier survival curve and log rank tests were performed using GraphPad Prism v7.06 software (GraphPad Software, Inc., San Diego, CA, USA).

### Statistical analysis

Statistical analysis was performed using a one/two-way analysis of variance with the means of three independent experiments unless otherwise specified. GraphPad Prism (GraphPad Software, San Diego, CA, USA) software package was used to calculate *P*-values. Results are means ± S.D. *: 0.01 ≤ *P* < 0.05; **: 0.001 ≤ *P* < 0.01; ***: *P* < 0.001.

## Results

### eEF2K promotes the translation of PD-L1

In some types of cells, the expression of PD-L1 is increased by interferon-γ (IFNγ [23]). IFNγ did indeed increase the levels of the PD-L1 protein in prostate cancer PC3 (human prostate cancer) cells (Fig. 1A, data quantified in Fig. 1B). Since no highly specific or potent small molecule inhibitor of eEF2K is yet available (even at high doses, the published compound A484954 [24] only weakly inhibited the phosphorylation of eEF2; Fig. 1A), we employed genetic approaches to knock down or knock out the expression of eEF2K. We used CRISPR-Cas9 genome editing to disrupt the *EEF2K* gene in PC3 cells (Fig. S1); as expected, since eEF2K is the only kinase that phosphorylates the regulatory site, Thr56, in eEF2 [25], no phosphorylation of eEF2 at this site was seen in eEF2K-KO PC3 cells (Fig. 1A). eEF2K was also undetectable in the knockout (KO) cells. eEF2K-KO showed lower levels of PD-L1 than the corresponding control cells (Fig. 1A,B). IFNγ effectively increased PD-L1 protein levels in the wild-type (WT) PC3 cells but not in eEF2K-KO cells. Thus, eEF2K positively regulates the expression of the PD-L1 protein in PC3 cells.

**Figure 1.**
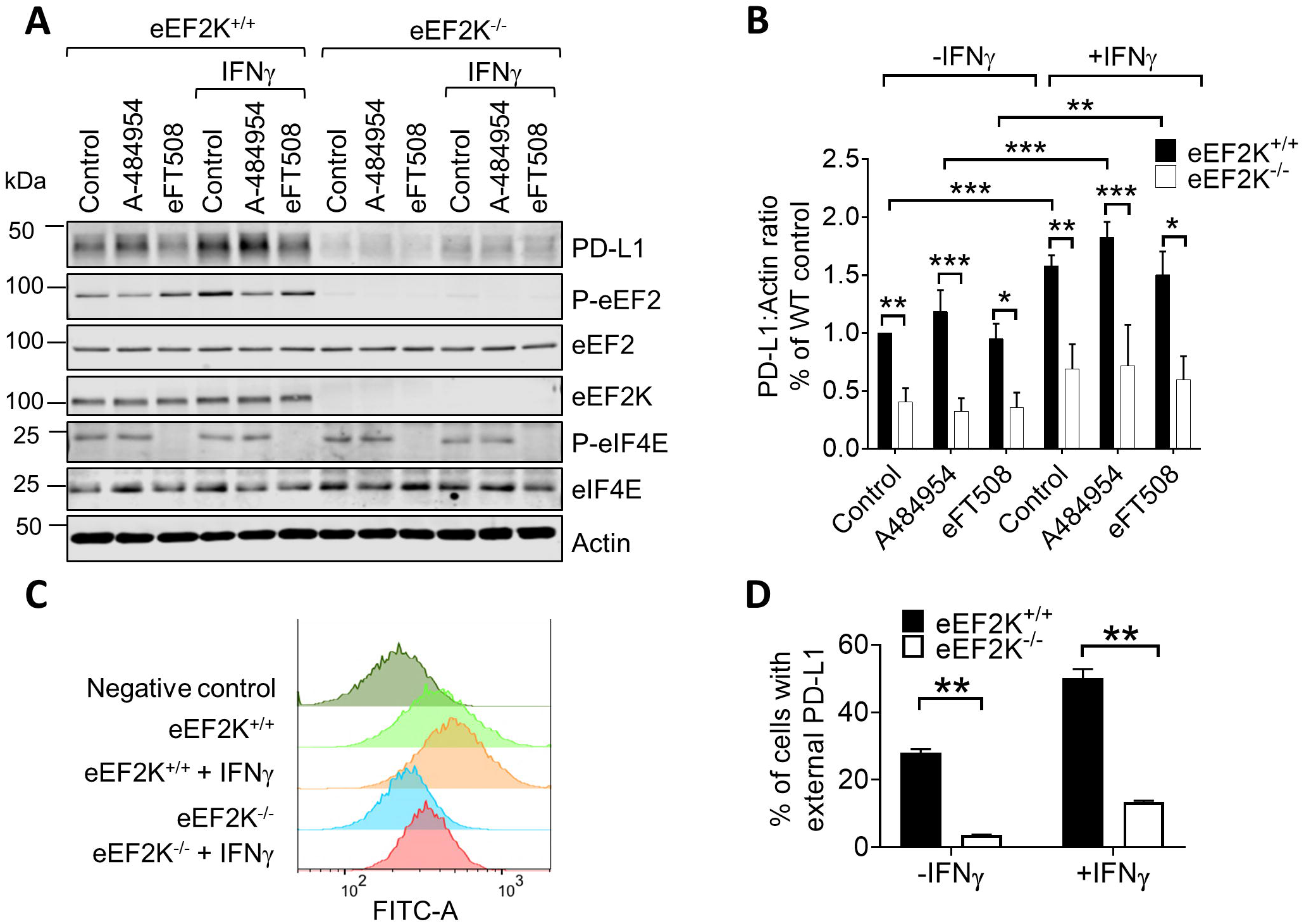
eEF2K upregulates PD-L1 protein expression in PC3 cells. **(A)** Cells were cultured in the presence or absence of 100 μM A-484954, 100 nM eFT508 and/or 20 ng/ml IFNγ for 24 h. The levels of indicated phospho-(P-) or total proteins were analysed by Western blotting. **(B)** Quantification of PD-L1 levels in **A** (*n* = 3). **(C)** Cells were treated with 20 ng/ml IFNγ for 24 h before subjected to flow cytometry analysis of cell surface PD-L1 protein expression. Negative control indicates cells “stained” without primary antibodies. **(D**) Quantification of **C**. Results are presented as means ± S.D.; **: 0.001 ≤ *P* < 0.01; ***: *P* < 0.001 (two-way ANOVA).

A recent study reported that the MNK kinases, which phosphorylate another component of the protein synthesis machinery, eIF4E, modulate the expression of PD-L1 [26]. However, at least in PC3 cells, inhibition of the MNKs and thus of the phosphorylation of eIF4E using eFT508 [27] had no effect of PD-L1 protein levels (Fig. 1A).

The key feature for immune-mediated killing of cancer cells is the amount of PD-L1 at the cell surface, which we assessed by FACS analysis. Consistent with the data for total PD-L1, in PC3 cells, knocking out eEF2K markedly decreased the amounts of PD-L1 detected at the cell surface, both in control cells and those treated with IFNγ (Fig. 1C, data are quantified in Fig. 1D).

We also made use of previously-described A549 human pulmonary adenocarcinoma cells harbouring an IPTG-inducible shRNA against the *EEF2K* mRNA [25] (Suppl. Fig. S2). Two stress conditions which activate eEF2K, oxidative stress (H_2_O_2_) and acidosis (maintaining cells at pH 6.7, [15]) increased PD-L1 protein levels (Suppl. Fig. S2A) in wildtype A549 cells. In contrast, PD-L1 protein levels trended lower in eEF2K-KO A549 cells under all conditions tested and were significantly reduced in cells treated with H_2_O_2_ or subjected to acidosis (Fig. S2A, quantified in Fig. S2B). [Note that, upon activation, eEF2K is sometimes degraded (via the proteasome [19, 28]). Similarly, PD-L1 levels were also reduced in eEF2K-null MDA-MB-231 cells (Fig. S2C and D) under conditions where eEF2K is active (rapamycin treatment, H_2_O_2_ or acidosis). Conversely, ectopic expression of FLAG-tagged eEF2K induced the expression of PD-L1 in A549 cells (Fig. S2E), in line with positive regulation of the expression of PD-L1 by eEF2K. Knockdown of eEF2K had little or no effect on total PD-L1 mRNA levels in A549 cells (Fig. S2F).

### eEF2K promotes the association of the PD-L1 mRNA with polysomes

Given that the only known substrate for eEF2K is eEF2, the protein which mediates the translocation of the elongation phase of mRNA translation, it seemed possible the eEF2K affected the translation of the *PD-L1* mRNA. To assess this in PC3 cells, we resolved lysates from control or eEF2K-KO cells on sucrose density gradients to separate non-polysomal mRNA from those associated with ribosomes or translationally-active polysomes of various sizes which contain ribosomes actively translating mRNAs. Cells were treated with IFNγ prior to lysis, to increase the levels of P-eEF2 (see Fig. 1A). Knockout of eEF2K did not affect the overall proportion of ribosomes in polysomes (Fig. 2A and B) or the distribution of a ‘control’ mRNA, that encoding β_2_-microglobulin (Fig. 2E, quantified in Fig. 2F). In marked contrast, 30% less of the PD-L1 mRNA was found in active polysomes in samples from PC3 cells in which eEF2K had been knocked down (Fig. 2C and D), and much more was in fractions corresponding to inactive or poorly translated mRNAs. This could be because residual eEF2K and P-eEF2 in IPTG-treated A549 cells were enough to sustain basal levels of *PD-L1* mRNA.

**Figure 2.**
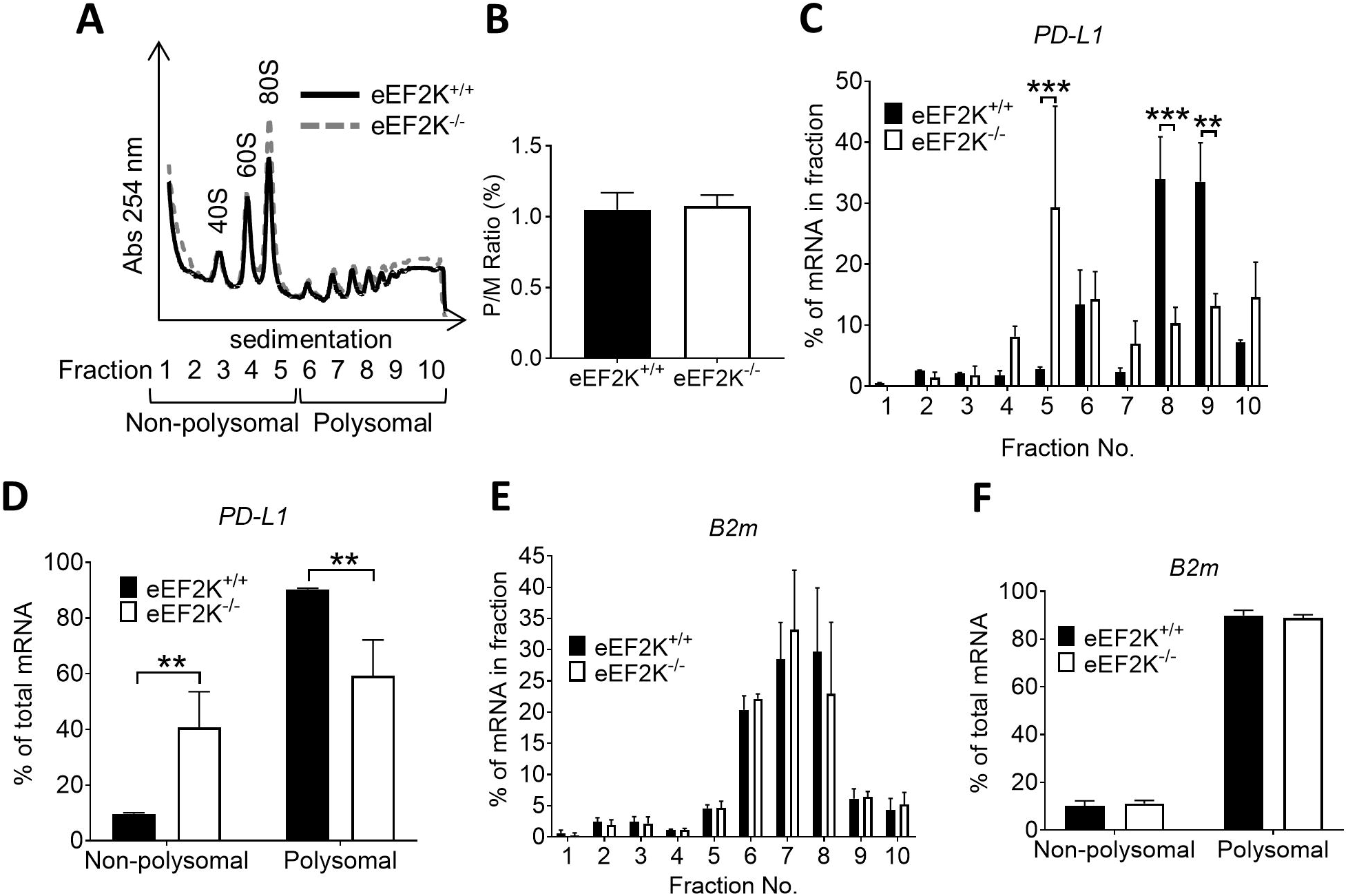
Ablation of eEF2K shifts *PD-L1* mRNAs from polysomal to non/subpolysomal fractions in PC3 cells. **(A)** Cells were treated with 20 ng/ml IFNγ for 24 h before subjected to sucrose density gradient analysis; representative images from three-independent experiments are shown. **(B)** The P/M (polysomal/non-polysomal) ratio from **A** was calculated by integrating the areas under the polysomal and non-polysomal fractions. qPCR was performed using specific primers for human **(C** and **D)** PD-L1 or **(E** and **F)** B2M. Positions of ribosomal/polysomal species are indicated. Results are presented as means ± S.D.; *0.01 ≤ P < 0.05; **: 0.001 ≤ *P* < 0.01; ***: *P* < 0.001 (two-way ANOVA).

Knockdown of eEF2K in A549 cells cultured with IFNγ did not affect the overall proportion of ribosomes in polyribosomes (Fig. S3A), but RT-qPCR analysis of the fractions revealed that, while a substantial proportion of the *PD-L1* mRNA was actively translated in control cells, there was almost no *PD-L1* mRNA was in polysomal fractions in eEF2K-knockdown cells (Fig. S3B). Instead, the *PD-L1* mRNA was found in the non-polysomal regions of the gradient, indicating that it was barely translated under glucose starvation. Thus, eEF2K promotes the association of the *PD-L1* mRNA with active polyribosomes in both PC3 and A549 cells, consistent with the observed changes in PD-L1 protein levels.

We noted that total *PD-L1* mRNA levels were decreased in eEF2K-KO PC3 cells under basal, rapamycin or oxidative or acidotic stresses or following IFNγ treatment (Fig. 3A and B). The *PD-L1* mRNA is reported to have a short half-life [29]; it was therefore plausible that the untranslated *PD-L1* messages in eEF2K-KO cells were rapidly degraded in PC3 cells. Indeed, when transcription of new mRNAs was blocked by actinomycin D, we observed that *PD-L1* mRNA levels decreased faster in eEF2K-null PC3 cells under both basal and IFNγ-treated conditions compared to the WT cells (Fig. 3C and D), indicating that the *PD-L1* the mRNA is less stable (has a shorter half-life) when eEF2K has been genetically deleted. This presumably reflects the decreased translation and, as a consequence, the impaired stability, of this mRNA under those conditions, explaining the differences in total *PD-L1* mRNA between control of eEF2K-KO PC3 cells. However, we cannot formally exclude that eEF2K affects *PD-L1* mRNA levels through additional mechanisms, which may also help explain the differences between A549 and PC3 in terms of the effect of knocking down or knocking out eEF2K on *PD-L1* mRNA levels.

**Figure 3.**
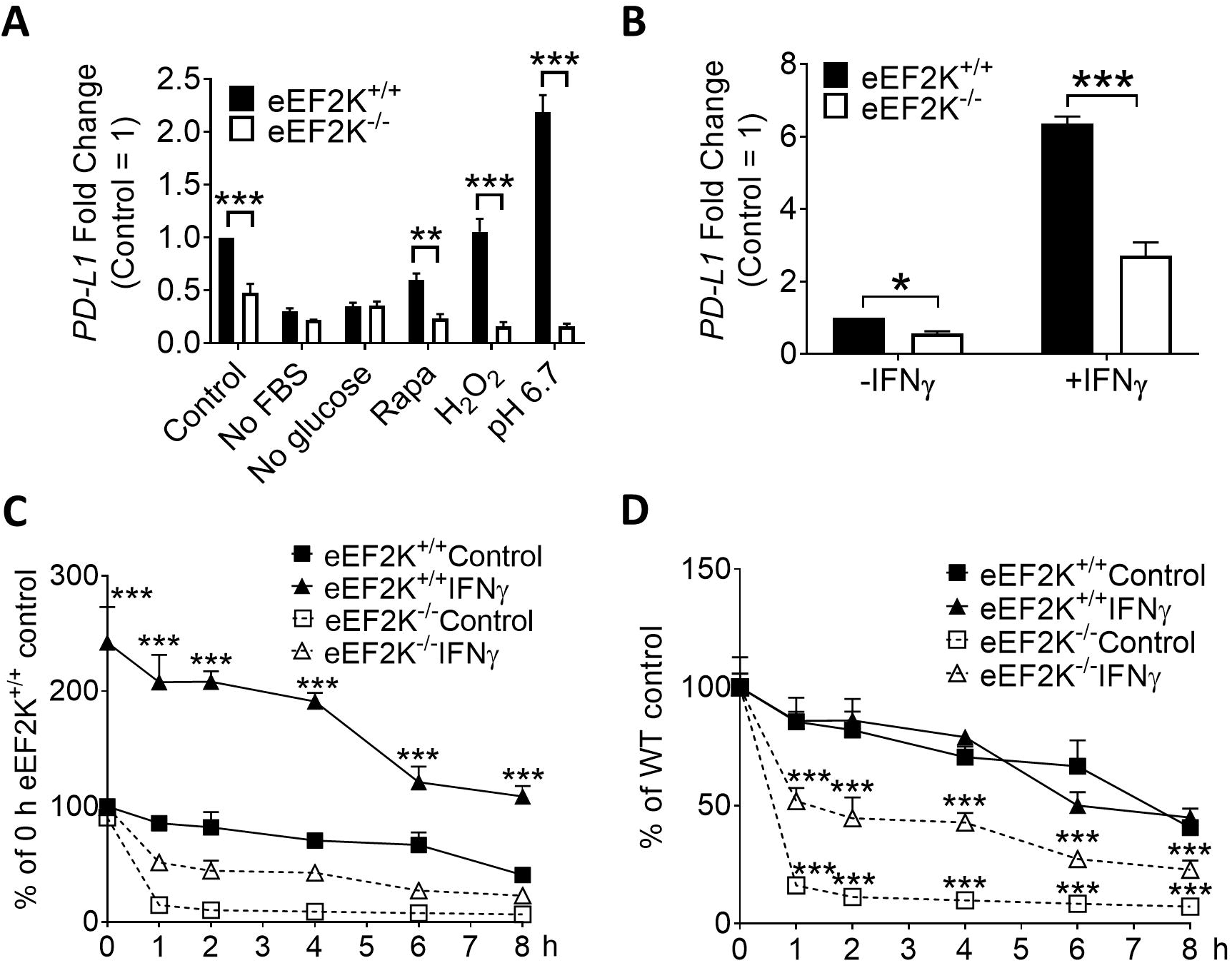
*PD-L1* mRNAs are more rapidly degraded in eEF2K-null PC3 cells. **(A)** Cells were cultured in in growth medium (control), or medium without FBS (no FBS), or medium without glucose (no glucose), or in growth medium with the presence of 200 nM rapamycin or 3 mM H_2_O_2_, or medium buffered to pH 6.7; for 24 h. **(B)** Cells were cultured in growth medium with or without 20 ng/ml IFNγ for 24 h. **(C** and **D)** Cells were cultured in growth medium with or without 20 ng/ml IFNγ for 16 h, before the addition of 2 μg/ml actinomycin D. Cells were further incubated for the indicated periods of time. (**A**-**D)**: qPCR was performed to monitor the levels of *PD-L1* mRNA. Results are presented as means ± S.D. (*n* =3); **: 0.001 ≤ *P* < 0.01; ***: *P* < 0.001 (two-way ANOVA).

### eEF2K enhances translation of the PD-L1 mRNA by bypassing translational repression mediated by an upstream open reading frame (uORF)

Control of the translation of specific mRNAs is often mediated through features of their 5’-untranslated region (5’-UTR, [30]). Regulatory elements such as upstream open reading-frames (uORFs) can modulate downstream reinitiation at the start of the main coding region of an mRNA; in general, translation of the uORF represses expression of the protein encoded by the main open reading-frame (ORF), although other scenarios may arise. The human *PD-L1* mRNA contains a single uORF which starts with CUG and partially overlaps the main *PD-L1* ORF (Fig. 4A); this overlap means that translation of this uORF would prevent expression of PD-L1 protein as ribosomes would terminate translation of this uORF after the start of the main PD-L1 ORF. (In contrast, the mouse *Pd-l1* mRNA contains three possible uORFs starting with either an AUG or a non-canonical CUG (uCUG) [26]). We recently reported that active eEF2K, and thus slower elongation, favors the selection of the canonical AUG as a start codon by augmenting the ability of ribosomes to distinguish between AUG and near-cognate start codons [28].

**Figure 4.**
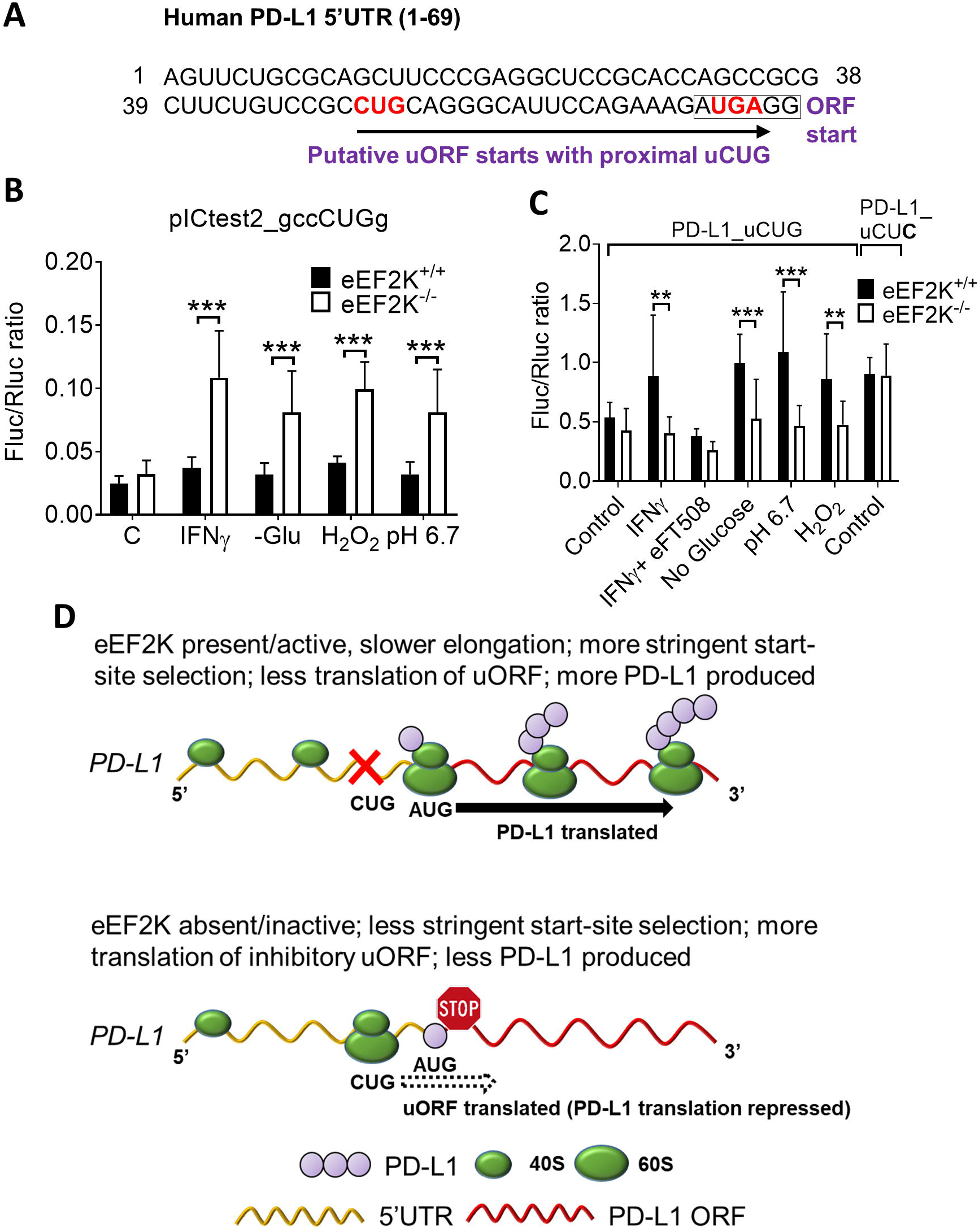
*PD-L1* translation is upregulated by eEF2K through a bypass of uORF-mediated translational repression in PC3 cells. **(A)** The sequence of the human *PD-L1* 5’UTR which contains a conserved putative uORF in mouse. Start (CUG) and stop (UGA) codons are in bold. **(B)** Cells were transfected with pICtest2 vectors encoding Fluc with CUG as a start codon. **(C)** Cells were transfected with PD-L1_uCUG or PD-L1_uCUC. **B** (*n* = 8) and **C** (*n* = 12): 24 h after transfection, cells were cultured in growth medium (control), treated with 20 ng/ml IFNγ (with or without 100 nM eFT508), or in medium without glucose (no glucose/-Glu), or in growth medium with the presence of 3 mM H_2_O_2_ or in pH 6.7-buffered medium; for 16 h. Fluc activity was then measured. Results are presented as ratios between Fluc and Rluc (means ± S.D.); **: 0.001 ≤ *P* < 0.01; ***: *P* < 0.001 (Student’s *t*-test). **(D)** Schematic representation of the mechanism by which eEF2K favors the selection of AUG over CUG as a start codon. C: control.

We have previously developed “pICtest2” dual-luciferase reporters with CUG as the start codon [17]. These constructs were created encoding Fluc as well as a cistron encoding *Renilla* luciferase as an internal control. eEF2K-KO PC3 cells were transfected with vectors expressing active Fluc from CUG or AUG as the start codon. Following IFNγ treatment or under various stress conditions (glucose starvation, oxidative and acidotic stress), eEF2K-KO PC3 cells exhibited a 2-3 fold higher levels of Fluc activity from the vector which has CUG as the start codon than WT (eEF2K+/+) cells (Fig. 4B). In contrast, the presence or absence of eEF2k did not affect the Fluc levels observed with the vector which has AUG as the start of the Fluc cistron (Fig. S4A) which is in line with our earlier data [28].

Next, we created a pICtest2 dual-luciferase reporter containing the full-length human *PD-L1* 5’-UTR (Fig. 4C) and transfected it into WT and eEF2K-KO PC3 cells. IFNγ treatment or stress conditions (glucose starvation, oxidative and acidotic stress) further enhanced Fluc activity in WT but not in eEF2K-KO or cells treated with IFNγ plus eFT508 [26] (Fig. 4C). In contrast, when transfected with a mutant vector where the uCUG was mutated to CUC (which is not used as a start codon; PD-L1_uCUC construct, see Fig. S4B) [26, 31], WT and eEF2K-null PC3 cells exhibited similar levels of Fluc activity (data not shown and Fig. 4C). Notably, consistent with previous observations [26], disrupting the uCUG increased the reporter activity by approximately 80% in both WT and eEF2K-KO control cells indicating that it does indeed impair translation of the main downstream ORF (Fig. 4C). These data clearly indicate that active eEF2K favors higher rates of initiation and translation of the main PD-L1 ORF, by slowing down elongation, enhancing the stringency of start site selection and thus reducing translation of the uORF (as depicted in the cartoon in Fig. 4D). We cannot absolutely rule out that there may be additional explanations for the ability of eEF2K to enhance the synthesis of PD-L1.

### Ablation of eEF2K enhances the ability of immune cell killing

We next sought to investigate whether the down-regulation of PD-L1 expression upon eEF2K ablation promote the capacity of immune cells to induce cancer cell death by performing immune killing assays. PD-1/PD-L1 inhibitor 3 was used as a positive control [32]. To correct for any changes in total cell number due to effects of proliferation or survival caused by knock-down of eEF2K, assay periods were kept short and, crucially, for any given condition data are expressed as number of surviving cells with added NK cells / total remaining cells without NK cells. Notably, both PC3 and A549 cells expressing sh-eEF2K, which express lower levels of PD-L1 (Fig. 5A and Fig. S2E, respectively) were more susceptible to immune killing by NK-92 cells than cells expressing sh-NC (Fig. 5B and Fig. S5; data presented as % cell survival ± NK cells to allow for any differences in total cell number). In terms of cancer cell survival, eEF2K only comes into play under conditions such as low nutrient availability [6], acidosis [15] or hypoxia [25] (reviewed in [8, 33]; Fig. 5B and Fig. S2A-D). Conversely, over-expression of FLAG-eEF2K, which increased the levels of PD-L1 protein (Fig. 5A, Fig. S2C) alleviated the cytotoxic effect of NK-92 cells towards PC3 and A549 cells (Fig. 5B, Fig. S2G and S5).

**Figure 5.**
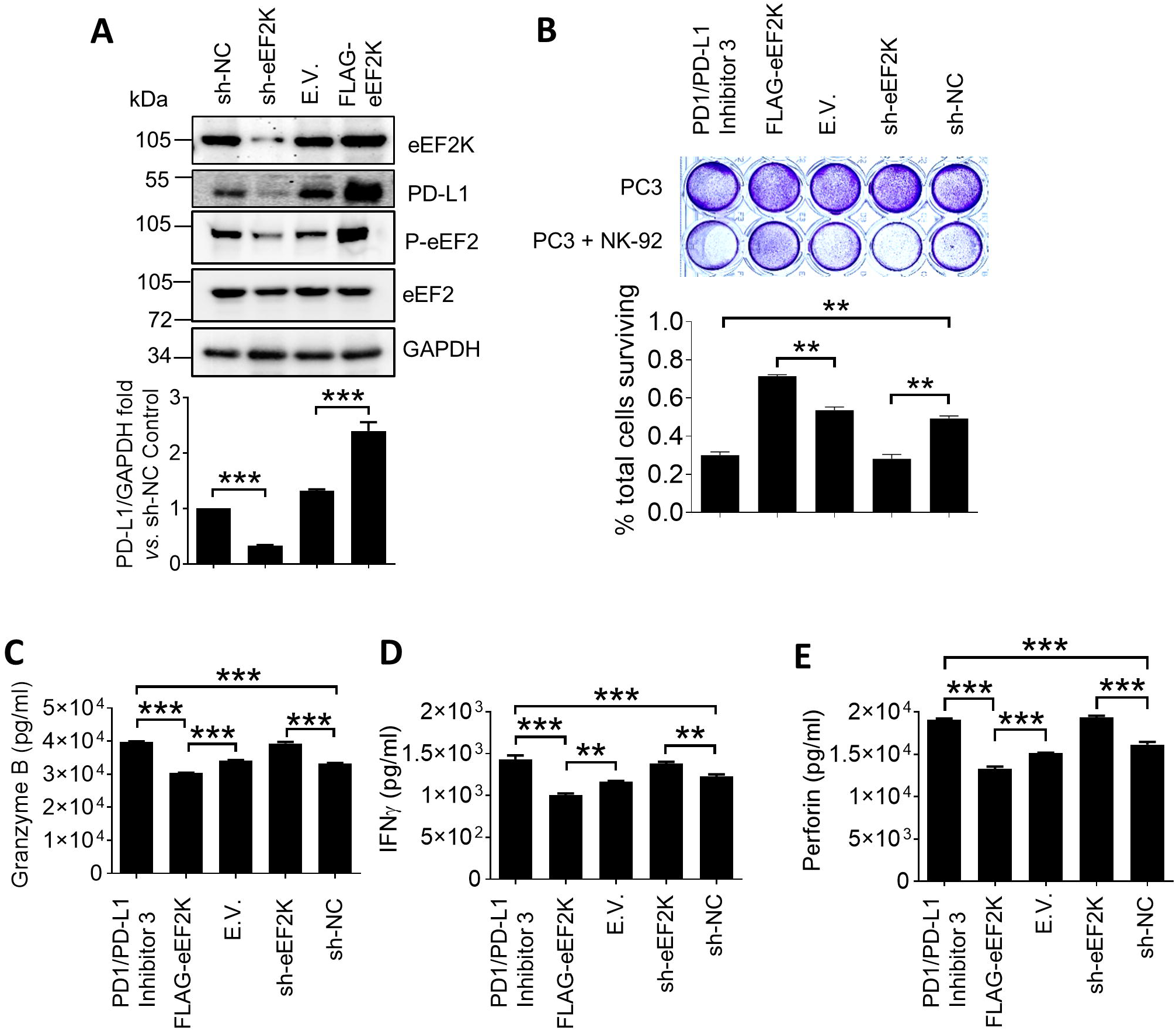
eEF2K-null PC3 cells are more susceptible to immune killing. PC3 cells were infected with sh-NC or sh-eEF2K, or transfected with an empty vector (E.V.) or FLAG-eEF2K. 48 h later, cells were co-cultured with NK-92 cells for another 3 h period of time, before being subjected to immunoblotting analysis **(A)** or crystal violet staining **(B)**. Cells treated with PD-1/PD-L1 inhibitor 3 (10 μM) were used as a positive control. Data are graphed as surviving cells ± NK cells, to compensate for possible differences in total cell number arising from altered eEF2K levels. Culture supernatants from **B** were also harvested after 12 h and analyzed by ELISA for **(C)** granzyme B, **(D)** IFN-γ and **(E)** perforin. Results are shown as means ± S.D., *n* = 3. *0.01 ≤ P < 0.05, ** 0.001 ≤ P < 0.01 (one-way ANOVA). For simplicity, not all instances of statistical significance are shown.

Consistent with these findings, as assessed by ELISA, levels of granzyme B, IFNγ and perforin in culture media from A549 cells over-expressing FLAG-eEF2K that had been co-cultured with NK-92 cells were lower compared to medium from cells transfected with an empty vector (Fig. 5C-E). Conversely, levels of these cytokines from A549 cells expressing sh-eEF2K and co-cultured with NK-92 cells were increased in comparison to the sh-NC expressing ones (Fig. 5C-E). These data further support the notion that over-expression of eEF2K in cancer cells blunts the immune response whereas eEF2K depletion enhances the susceptibility of cancer cells towards immune attack.

### Gene expression profiles of EEF2K and CD274 (PD-L1) in a database of prostate cancer patients

In addition, gene expression profiles retrieved from the TCGA (Fig. 6A) and Grasso (Fig. 6B) databases have demonstrated that the expression levels of *EEF2K* and *CD274*, which encodes PD-L1, are positively correlated in tumour samples obtained from prostate cancer patients [21, 22]. Notably, high levels of *EEF2K* expression were associated with decreased overall survival of prostate cancer patients (Fig. 6C). High levels of *CD274* expression were also associated with decreased progression-free survival of lung cancer patients (Fig. 6D). In contrast, low levels of *EEF2K* expression were associated with lower progression-free survival of lung cancer patients (data not shown). This is in line with the fact that the role of eEF2K in cancer development is complex as revealed by apparently conflicting reports, which suggest eEF2K may promote or impair tumour growth or survival. For example, we recently reported that depletion of eEF2K promotes growth of A549 xenografts in mice [34]. In direct contrast, eEF2K is also highly expressed in other cancers (see, e.g., [35, 36] and is also activated under conditions (such as nutrient depletion or hypoxia [8]) which occur within solid tumours; this will permit the enhanced expression of PD-L1 increasing the resistance of the tumour cells to immune attack.

**Figure 6.**
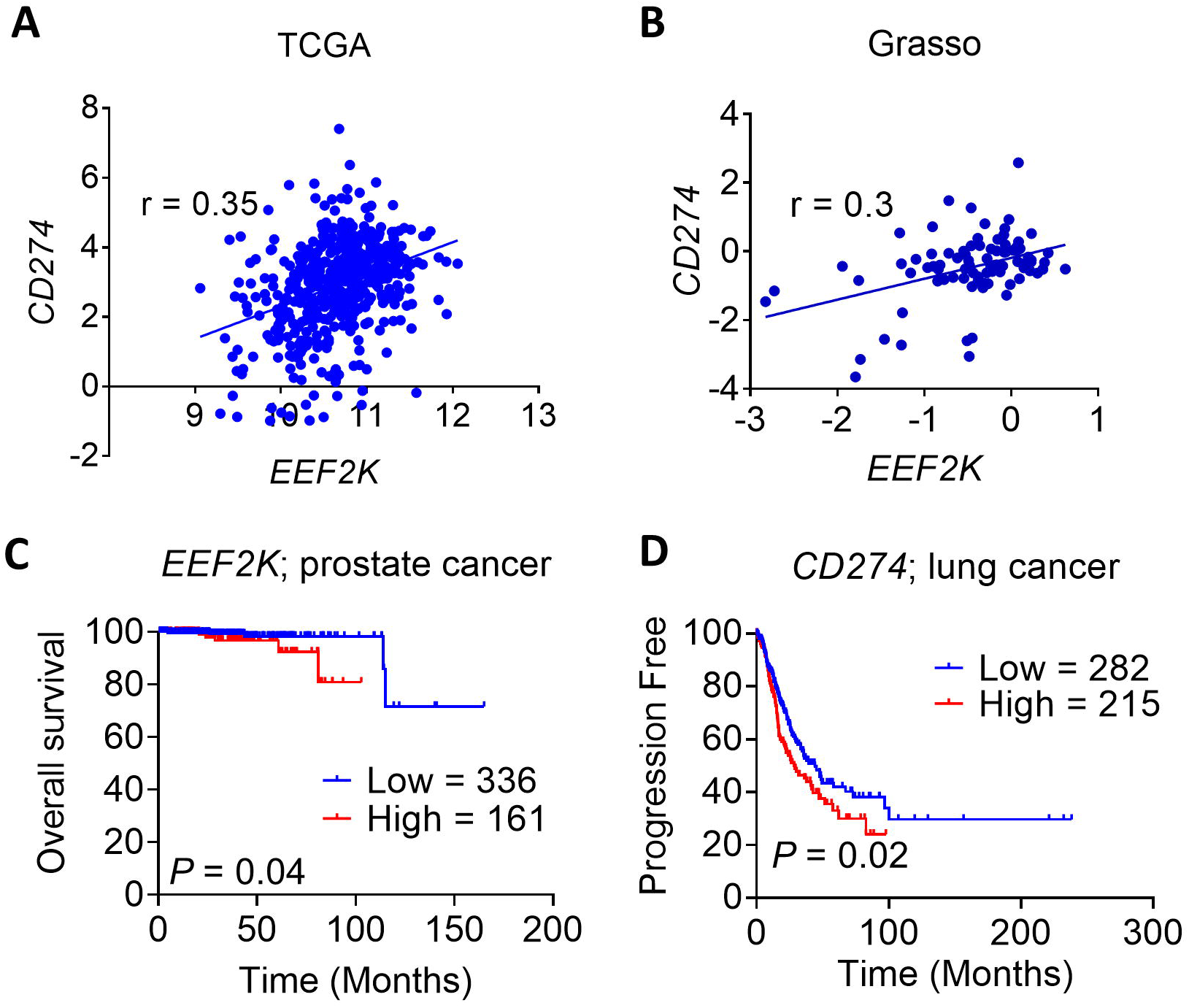
High levels of *EEF2K* positively correlate with enhanced *CD274* gene expression and poor survival rates in cancer patients. **(A)** Pearson’s correlation analysis between the expression of *EEF2K* and *CD274* in TCGA (prostate cancer) and, (**B)** Grasso datasets (prostate cancer). Kaplan-Meier plots showing overall survival in **(C)** prostate cancer and **(D)** progression-free survival in lung adenocarcinoma with low/high *EEF2K* **(C)** or *CD274* **(D)**, respectively. Data were obtained from TCGA datasets. Patients were stratified into two groups, according to the level of mRNA expression of *EEF2K* or CD274. Data were statistically analysed using a two-sided log-rank test.

## Discussion

Here we show that eEF2K positively regulates the levels of the major immune checkpoint protein, PD-L1 in three different types of cancer cells (A549, PC3 and MDA-MB-231) and provide evidence that this occurs at the level of the translation of its mRNA. This adds to the growing evidence that eEF2K helps to promote tumour growth and metastasis, providing further impetus to the concept that eEF2K (a non-essential and atypical protein kinase) is an attractive target for the therapy of at least some types of solid tumours [8-10, 37]. eEF2K is activated under conditions which occur within tumours, such as nutrient depletion [6] or hypoxia [25]; this will also facilitate the production of PD-L1 and thus enhance the tumour’s resistance to immune attack. Further work, using e.g., murine models, will be needed to explore the role of eEF2K in protecting tumour cells from immune surveillance *in vivo*; as our cancer cells in which eEF2K has been depleted or knocked out are of human origin, we are currently unable to perform such studies which require immunocompetent mice.

Inhibition of mTORC1 and its substrate S6K, both of which are negative regulators of eEF2K [38, 39], has also been shown to promote the synthesis of PD-L1 in lung cancer cells [40], although in that case this was attributed to effects on the stability of the PD-L1 protein. Furthermore, the mTOR inhibitor everolimus also increases PD-L1 expression in renal cell carcinoma lines [41], an effect which, at least in part, might involve the activation of eEF2K. In contrast, in several mutant EGFR (EGFR^L858R/T790M^) and KRas (KRas^LA2^ and KRas^G12D^) murine lung cancer models, active AKT-mTOR signaling actually increases PD-L1 expression [42]. Therefore, the impact of particular signalling pathways such as mTORC1 (and thus upstream PI 3-kinase signalling) on PD-L1 expression differs substantially between types of cancers or cancer cells [40-42].

While two earlier studies reported direct or indirect translational control of the regulation of PD-L1 protein levels, our findings are quite distinct, since our data reveal direct control of the translation of the *PD-L1* via control of translation elongation. Furthermore, our data provide the first example of control of translation of a uORF-containing mRNA via regulation of the translation elongation machinery. Thus, while eEF2K inhibits general protein synthesis, it can actually enhance the synthesis of PD-L1; in fact, there are now several published examples where eEF2K stimulates the translation of specific mRNAs [9, 11, 18]. Given that uORFs with a non-canonical CUG start codon appear to exist in many mRNAs [13], eEF2K may control the translation of certain other mRNAs in a similar way. Furthermore, there is a very well-known system whereby a phosphorylation event on another translation factor, eIF2, inhibits overall translation but actually promotes translation of some mRNAs [43]. Thus, two different mechanisms involving inhibitory phosphorylation of an initiation factor (eIF2) or elongation factor (eEF2), which generally operate to slow down translation, can actually promote translation of specific mRNAs, by virtue of features of their uORFs. Since eEF2K and the kinases that phosphorylate eIF2 are activated under different circumstances [3, 43], these mechanisms are poised to modulate the synthesis of specific proteins under distinct conditions.

In the study by Cerezo *et al*. [44], in melanoma, the eIF4F translation initiation factor complex was shown to positively regulate the translation of the mRNA encoding the transcription factor STAT1 and thus the induction of the *PD-L1* mRNA in response to IFNγ. Another study, on KRAS-driven liver cancer [26], also provided evidence that PD-L1 expression is controlled through eIF4F, but in this case through its direct control of the translation of the mRNA for PD-L1 through the phosphorylation of another component of that complex, eIF4E. We did test the same MNK inhibitor as Xu *et al*. [26], eFT508 [27], but saw no significant effect on PD-L1 protein levels in PC3 (Fig. 1A) or A549 (data not shown) cells. It thus appears that control of PD-L1 expression by the MNKs is restricted to some kinds of (cancer) cells.

An important implication of our data, together with other recent findings [26, 44] is that the translational regulatory networks somehow steer events to aid cancer cells to evade immunosurveillance by increasing the synthesis of immune checkpoints, and thus promote tumour cell migration and metastasis. In particular, the previously reported *PD-L1* 5’-UTR serves to allow cancer cells utilise regulators of mRNA translation, e.g. eEF2K, to increase the levels of a “cloak of invisibility” – PD-L1 - and thereby hide themselves from immune attack. Specifically, the human *PD-L1* 5’-UTR contains a conserved uORF which starts with a non-canonical start codon (CUG). The uORF partially overlaps with and inhibits the translation of the main ORF encoding PD-L1 [26]. We previously showed that activation of eEF2K helps reduce initiation at non-canonical start sites, i.e., CUG or GUG [28]. Impaired elongation slows down ribosomes within the main ORF and thus allows more time for selection of the optimal choice – an AUG start codon. Thus, by reducing initiation at the inhibitory uORF, eEF2K can promote translation of the main ORF and thus the production of PD-L1, as observed in our experiments. Our findings identify for the first time a specific feature of an mRNA which confers positive control of its translation by eEF2K. Since a substantial number of other mRNAs possess uORFs, and some of them may also be subject to translational control by eEF2K, although this is likely depends on other features such as the location of the uORF within the 5’-UTR and the nature of its start codon and its immediate context.

To further support the notion that immune checkpoint is tightly regulated by translation elongation, the present study also demonstrates for the first time a positive correlation between *EEF2K* and *CD274* levels in prostate cancer patients. Aberrantly increased *EEF2K* levels in prostate or lung cancer patients also predict poor survival. Importantly, our data show that depletion of eEF2K enhances the susceptibility of cancer cells to NK-cell mediated toxicity. Unfortunately, we cannot use immunocompetent mouse models to test these effects in vivo, as the available eEF2K-KO cell lines are of human origin.

Taken together with the roles of eEF2K in promoting the survival of cancer cells under nutrient-deprived conditions [6, 35, 36], in cell migration and cancer cell metastasis [9], and in angiogenesis [10], our present data add further support to the conclusion that disabling eEF2K may be a valuable therapeutic approach to tackling established solid tumours, at least for some types of cancers. However, as is the case with targeting other processes such as autophagy and mTORC1 signaling [2, 45, 46], in some specific settings the inhibition of eEF2K may actually promote cancer-related processes, e.g., tumour initiation. Blocking eEF2K function may be more appropriate in later stages of tumour progression although this may also depend on the genetic and/or metabolic profile of the tumour. Our data further highlight the multi-faceted role played by eEF2K in regulating protein expression and cellular functions in cancer cells.

## Author Contributions

Y. Wu, J. Xie, X. Wang, J. Li and C. G. Proud designed the experiments. Y. Wu, J. Xie, X. Jin and R. V.Lenchine performed research. D. M. Fang and Z. D. Nassar also contributed data. L.M. Butler, J. Li X. Wang and C. G. Proud provided supervision. All authors interpreted and analysed the data. J. Xie, X. Wang and C. G. Proud wrote the manuscript.

## Acknowledgements

We gratefully acknowledge financial support from the South Australian Health & Medical Research Institute (SAHMRI), Australia. We thank Dr Rui Liu (Molecular Genetics Unit, St. Vincent’s Institute of Medical Research, Fitzroy, VIC 3053, Australia) for designing a cloning strategy for the PDL1-uCUG vector. We also thank Dr Randall Grose from the Australian Cancer Research Foundation (ACRF) Cellular and Imaging Cytometry Core Facility at SAHMRI for his assistance and expertise in FACS cell sorting. Patient survival rates are based upon data generated by the TCGA Research Network: https://www.cancer.gov/tcga.

## Abbreviations

5’-UTR: 5’-untranslated region
BD: Becton Dickinson
eEF2: eukaryotic elongation factor 2
eEF2K: eEF2 kinase
eIF2: eukaryotic translation initiation factor 2
FBS: fetal bovine serum
Fluc: *firefly* luciferase
IFNγ: interferon-γ
IPTG: isopropyl-β-D-thiogalactopyranoside
mTORC1: mechanistic target of rapamycin complex 1
NEB: New England Biolabs
ORF: open reading-frame
PBS: phosphate buffered saline
PD-1: programmed cell death protein 1
PD-L1: PD-ligand 1
SDS-PAGE: sodium dodecyl sulphate-polyacrylamide gel electrophoresis
sh-NC: lentiviral non-targeting control
shRNA: short hairpin RNA
uORF: upstream open reading-frame
WB: Western blot

