## Supplemental Figure Legends for "eEF2 kinase enhances the expression of PD-L1 by promoting the translation of its mRNA"

**Supplementary Figure legends**

**Figure S1. Sanger sequencing analysis of CRISPR-directed eEF2K knockout in PC3 cells. Related to Figure 1.** The data reveal that, as a result of the CRISPR-Cas9 genome editing, part of exon 2 was deleted from the *EEF2K* gene in these PC3 cells.

**Figure S2. eEF2K is responsible for the induction of PD-L1 expression in A549 and MDA-MB-231 cells. Related to Figure 2. (A)** 5 days prior to experiment, eEF2K expression had been knocked down by an inducible (by IPTG) shRNA. Cells were cultured in growth medium (control), or medium without FBS (no FBS), or medium without glucose (no glucose), or in growth medium with the presence of 200 nM rapamycin or 3 mM H_2_O_2_, or medium buffered to pH 6.7; for 24 h, before lysis and immunoblotting analysis. **(B)** Quantification of **A**. Results are shown as means ± S.D., *n* = 5. *0.01 ≤ P < 0.05 (one-way ANOVA). **(C)** WT or eEF2K-null MDA-MB-231 cells were cultured in growth medium (control), or medium without FBS (no FBS), or without glucose (no glucose), or in growth medium with 200 nM rapamycin or 3 mM H_2_O_2_, or medium buffered to pH 6.7,; for 24 h, before lysis and immunoblot analysis. **(D)** Quantification of data in **C**. Results are shown as means ± S.D., *n* = 3. **0.001 ≤ P < 0.01; ***: *P* < 0.001 (one-way ANOVA). **(E)** A549 cells were infected with sh-NC or sh-eEF2K, or transfected with an empty vector (E.V.) or FLAG-eEF2K. 48 h later, cells were co-cultured with NK-92 cells for another 8 h period-of-time, before being subjected to immunoblot analysis. **(F)** samples from A549 cells treated as in **A** were subjected to RT-qPCR analysis to assess levels of *PD-L1* mRNA.

**Figure S3. Knocking down eEF2K in A549 cells shifts *PD-L1* mRNA from polysomal to non/subpolysomal fractions. Related to Figure 2. (A)** Sucrose density gradient analysis of lysates cells treated with 20 ng/ml IFNγ. **(B)** qPCR analysis of human PD-L1 from fractions from experiments shown in **A**.

**Figure S4. Translation of Fluc with AUG as a start codon is not affected by eEF2K in PC3 cells. Related to Figure 4. (A)** Cells were transfected with pICtest2 vectors encoding Fluc with AUG as a start codon. **(B)** Sanger sequencing analysis of pICtest2 PD-L1_uCUG/uCUC constructs. 24 h after transfection, cells were cultured in growth medium (control), treated with 20 ng/ml IFNγ (with or without 100 nM eFT508), or in medium without glucose (no glucose/-Glu), or in growth medium with the presence of 3 mM H_2_O_2_ or in pH 6.7-buffered medium; for 16 h. Fluc activity was then measured. Results are presented as ratios between Fluc and Rluc (means ± S.D.).

**Figure S5.** **eEF2K-depleted cells are more susceptible to immune killing.** **Related to Figure 5.** **(A)** Crystal violet staining of A549 cells corresponding to the type of experiment shown in Fig. S2E. Cells treated with PD-1/PD-L1 inhibitor 3 (10 μM) were used as a positive control. Data are graphed as surviving cells ± NK cells, to compensate for possible differences in total cell number arising from altered eEF2K levels. Results are shown as means ± S.D., *n* = 3. *0.01 ≤ P < 0.05, **0.001 ≤ P < 0.01 (one-way ANOVA). **(B)** PC3 cells were infected with sh-NC or sh-eEF2K, or transfected with an empty vector (E.V.) or FLAG-eEF2K. After 48 h, cells were co-cultured with NK-92 cells for another 3 h before being imaged using a Cytation 5 multimode Reader. Scale bar, 1 mm. Cells treated with PD-1/PD-L1 inhibitor 3 (10 μM) were used as a positive control.
