## Supplementary figures and images for "eEF2 kinase enhances the expression of PD-L1 by promoting the translation of its mRNA"

### Supplemental Figure 1

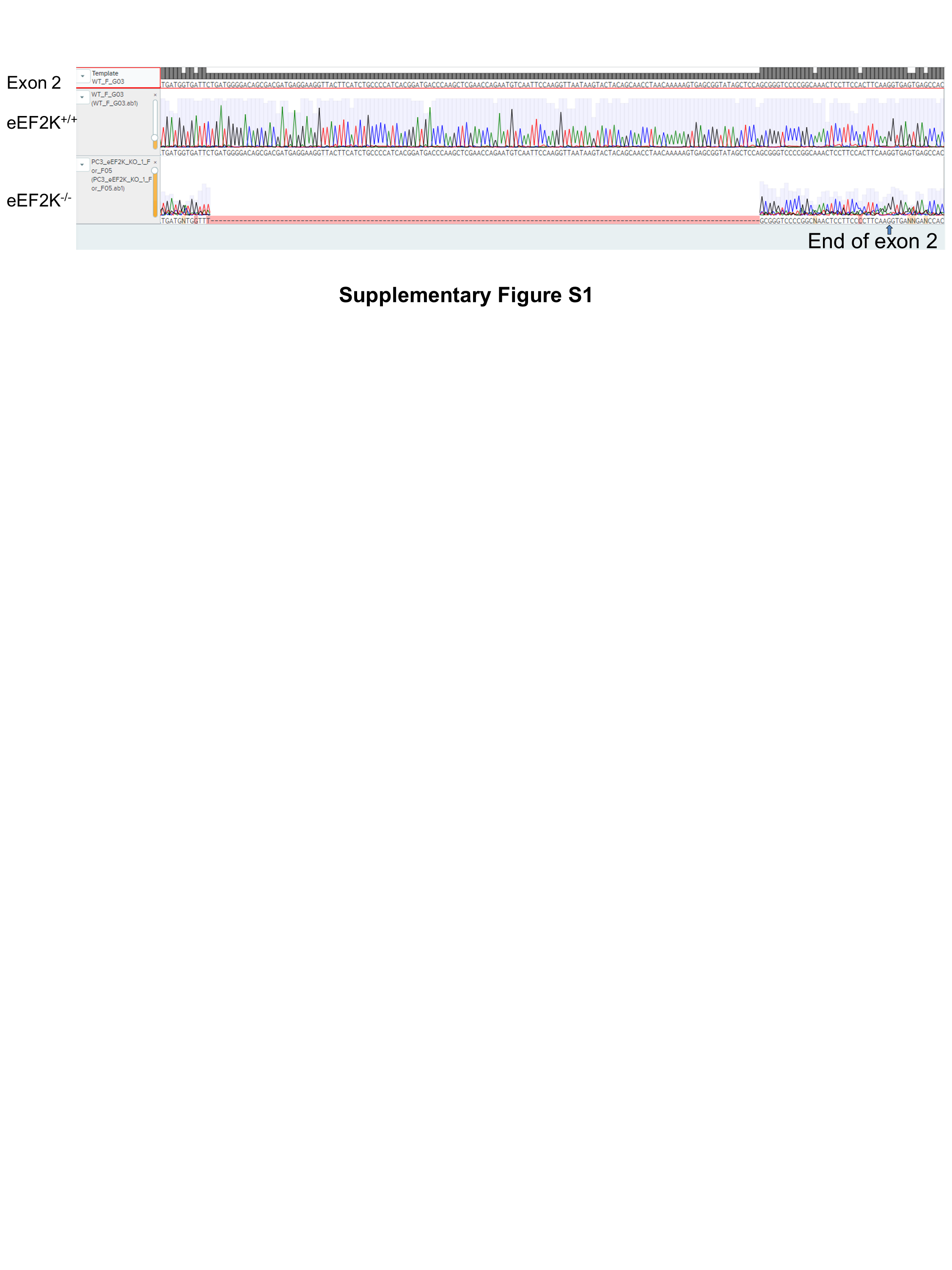

### Supplemental Figure 2

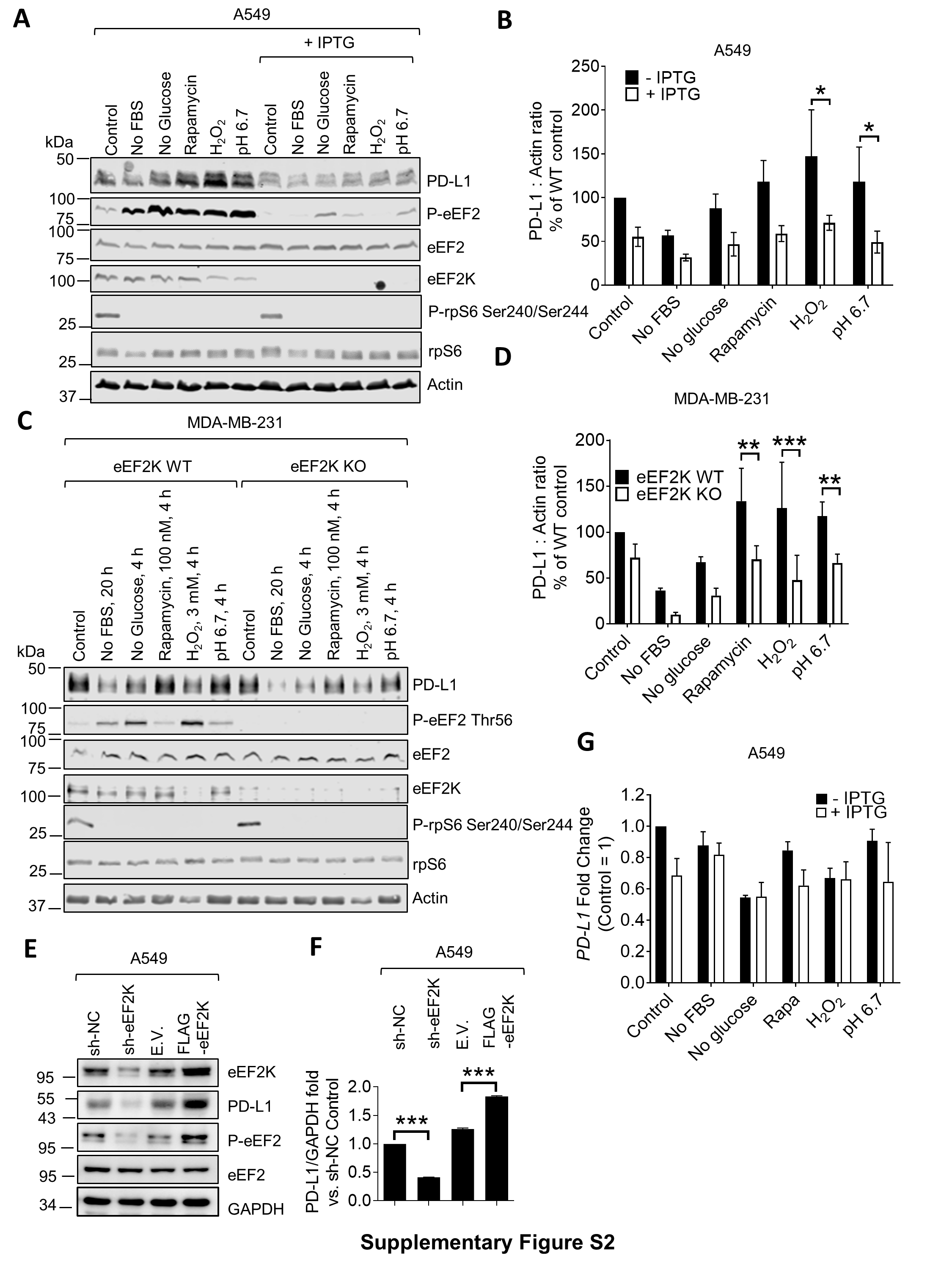

### Supplemental Figure 3

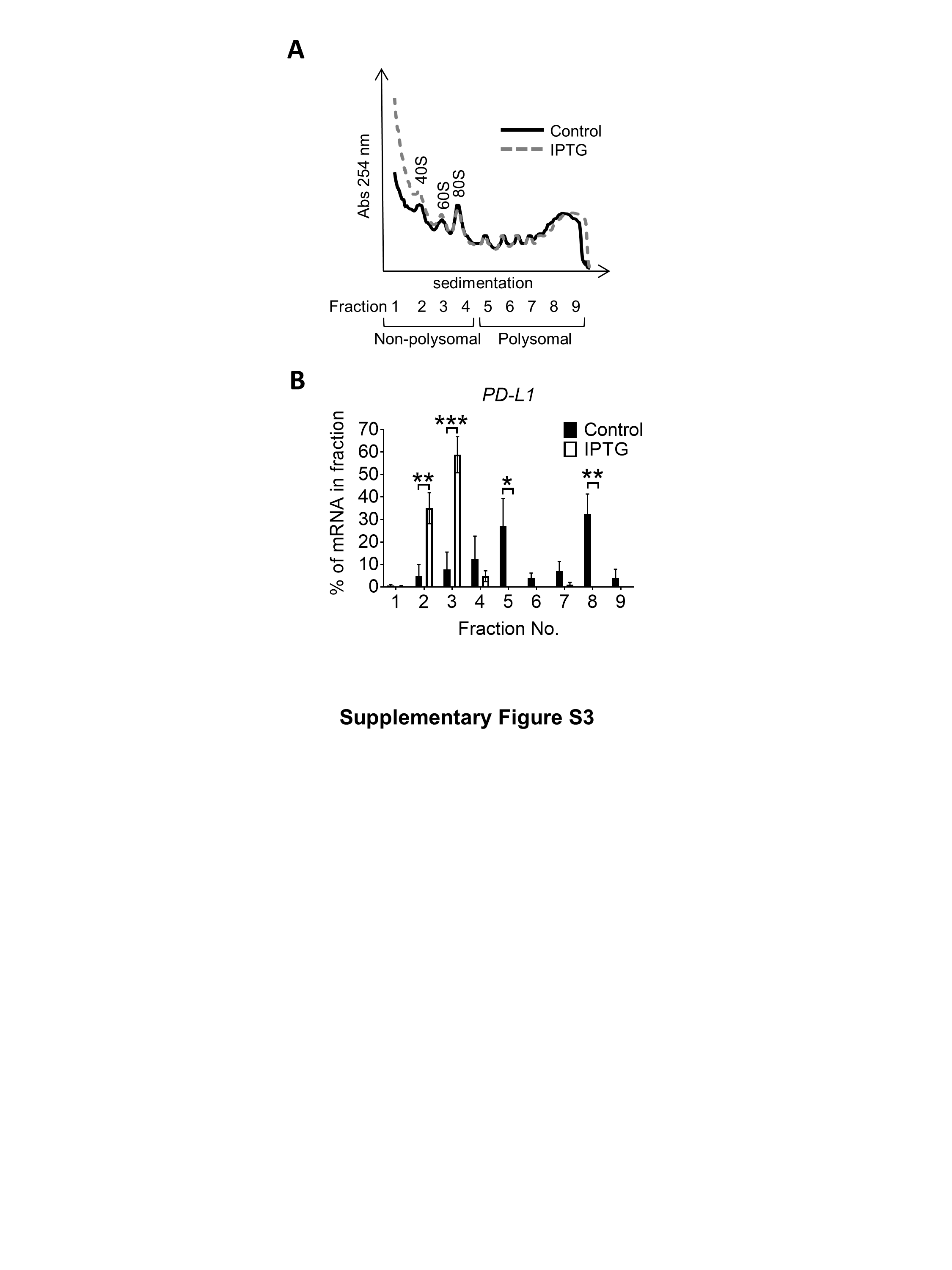

### Supplemental Figure 4

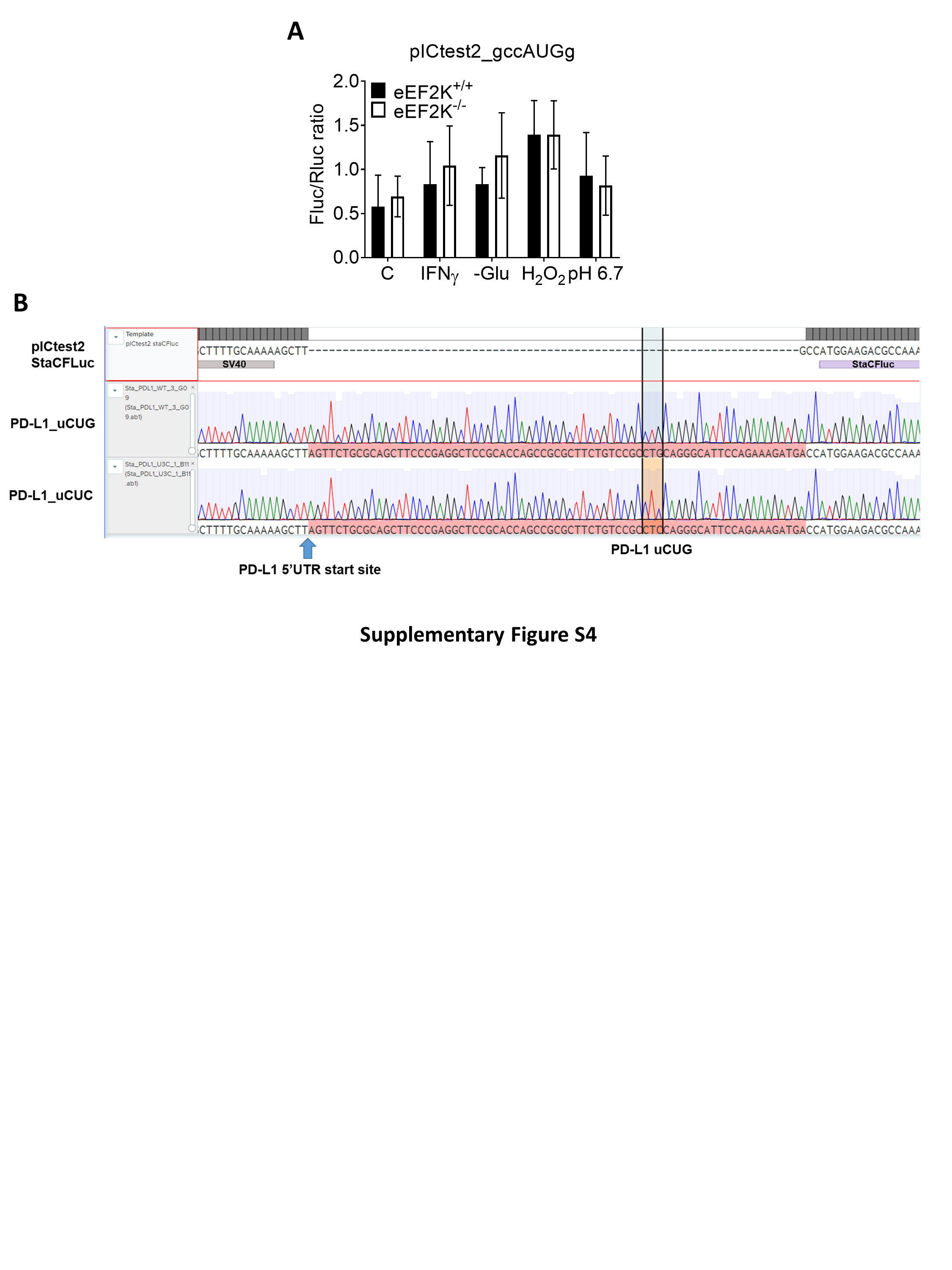

### Supplemental Figure 5

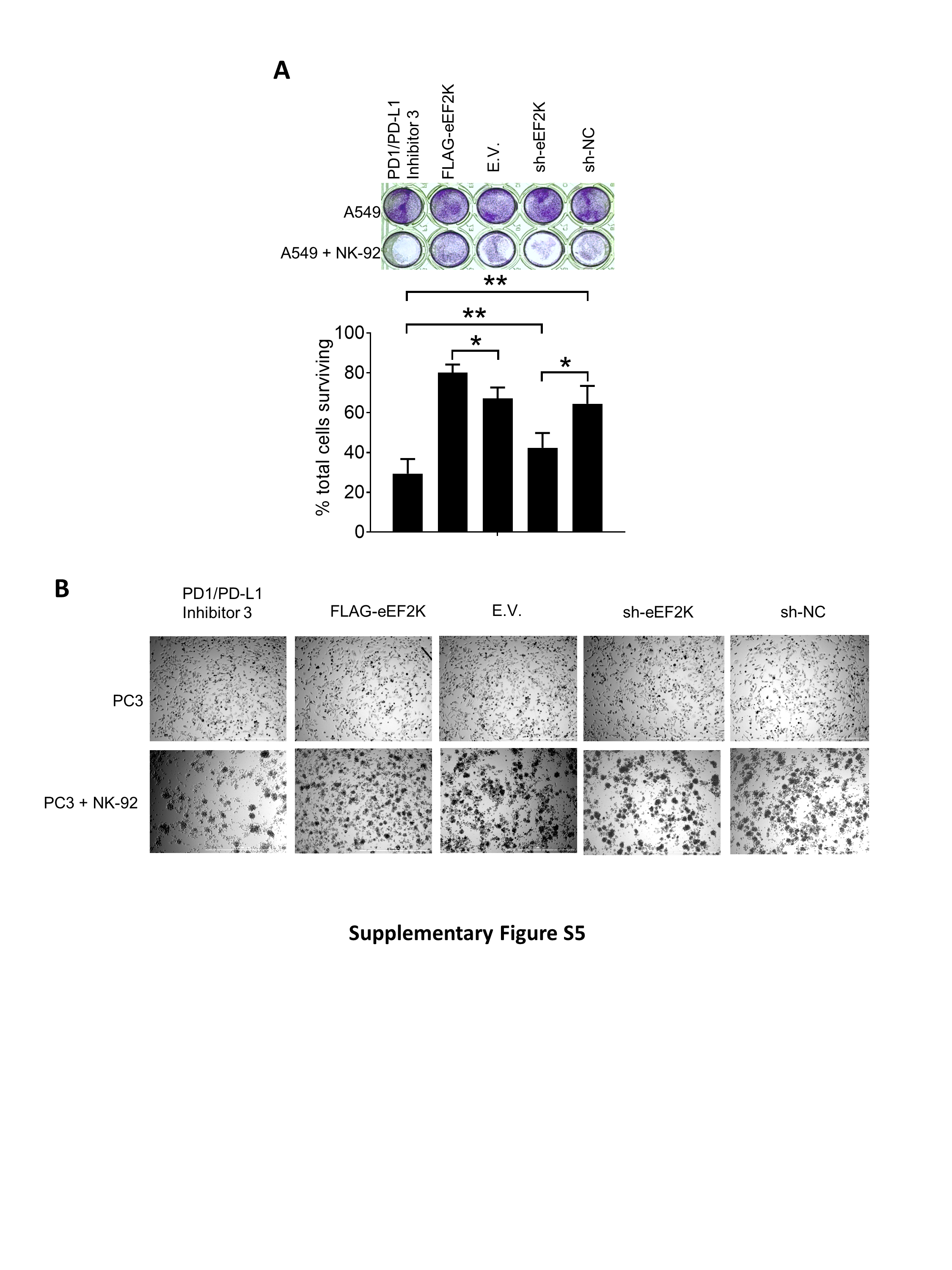
